# JUNB O-GlcNAcylation-mediated promoter accessibility of metabolic genes modulates distinct epithelial lineage in pulmonary fibrosis

**DOI:** 10.1101/2024.05.27.594700

**Authors:** Marie-Therese Bammert, Meshal Ansari, Leoni Haag, Zuhdi Ahmad, Victoria Schröder, Joseph Birch, Diana Santacruz, Werner Rust, Coralie Viollet, Benjamin Strobel, Alec Dick, Florian Gantner, Holger Schlüter, Fidel Ramirez, Muriel Lizé, Matthew J. Thomas, Huy Q. Le

## Abstract

Idiopathic pulmonary fibrosis (IPF) is a lethal disease with substantial unmet medical needs. While aberrant epithelial remodelling is a key factor in IPF progression, the molecular mechanisms behind this process remain elusive. Using a patient-derived 3D distal airway epithelial organoid model, we successfully recapitulate important IPF features, including the emergence of aberrant KRT5+/COL1A1+ basal cells and a metabolic shift towards increased O-linked β-N-acetylglucosamine (O-GlcNAc) levels. Consistent with this, single-cell analysis of accessible chromatin reveals an increased chromatin accessibility in these aberrant basal cells, particularly at JUNB motif-enriched promoter regions of metabolic genes. O-GlcNAcylation shapes JUNB function and promotes a pro-fibrotic response to chronic injury, leading to aberrant epithelial remodelling. Site-specific deletion of O-GlcNAcylation on JUNB attenuates the metaplastic differentiation of basal cells, thereby aiding in the restoration of the alveolar lineage. Together, these data establish a novel link between metabolic dysregulation, mediated by the O-GlcNAc-JUNB axis, and bronchiolization in IPF, offering new therapeutic strategies to treat this fatal disease.

## Introduction

The provision of cellular energy requirements is crucial in maintaining tissue homeostasis. This is achieved through cellular metabolism, which not only transforms nutrients into energy but also provides critical signals for other cellular activities, including proliferation, differentiation, immune response, and cytokine secretion^1–4^. These metabolic pathways are stringently regulated by a myriad of enzymes and signalling cascades, ensuring the fulfilment of the varied biological processes in accordance with their metabolic state. Any dysregulation within these processes can lead to injury and diseases^5^. One prominent metabolic rheostat - O-linked β-N-acetylglucosamine (O-GlcNAc) - is a unique nutrient- and stress-sensing glycosylation that functionally modifies a broad spectrum of proteins^6^. O-GlcNAc is transferred via the O-GlcNAc transferase (OGT) to serine and threonine residues of target proteins and is removed by the O-GlcNAcase (OGA), reflecting a unique dynamic property for post-translational glycosylation. OGT utilises uridine-diphosphate N-acetylglucosamine (UDP-GlcNAc) as a precursor, which originates from several nutrient-converting pathways, i.e. glycolysis, amino acid, or fatty acid metabolism. Regulatory proteins subject to O-GlcNAcylation are involved in the spatiotemporal regulation of a broad spectrum of cellular processes, from transcriptional regulation to stress response, to maintain tissue homeostasis. Dysregulation in O-GlcNAcylation is associated with certain pathologies including cancer, fibrosis, and ageing^7,8^. Idiopathic pulmonary fibrosis (IPF) is a rare, devastating disease, characterised by the excessive accumulation of extracellular matrix, leading to distortion of the alveoli and increased stiffness of the lung together with the loss of gas-exchange surface^9^. IPF patients have a poor prognosis, with an estimated survival time of two to three years post-diagnosis. While the causes of IPF remain poorly understood, it becomes evident that aberrant epithelial repair plays a major role in the progression of the disease^10,11^. Recent findings revealed an epithelial subpopulation of aberrant basal cells, marked by KRT5+/KRT17+/COL1A1+, promoting the pro-fibrotic cell fates and bronchiolization process in the lung^12,13^. However, the underlying mechanisms shaping this disease phenotype and the fibrotic cell fate of lung epithelial progenitors/stem cells remain elusive.

Harnessing a patient-derived 3D distal airway epithelial organoid model, we found that metabolic dysregulation, specifically through aberrant O-GlcNAc, determines the pro-fibrotic cell fate in IPF. This dysregulation shapes the hallmarks of IPF including expression of pro-fibrotic markers, increased proliferative capacity and presence of metaplastic aberrant basal cells by JUNB-O-GlcNAc. By inhibiting O-GlcNAcylated JUNB, we successfully mitigated these important fibrotic characteristics. This discovery illuminates a novel path in which dysregulated metabolism translates into cell fate decisions. Furthermore, it suggests a new therapeutic approach for IPF by targeting aberrant O-GlcNAc modifications on disease-related factors, offering a promising avenue for the treatment of IPF and other age-related diseases.

## Results

### Patient-derived distal airway organoids recapitulate IPF phenotype

We revisited published single-cell RNA sequencing (scRNA-seq) data from the lungs of IPF patients^14^, with a focus on aberrant basal cells (Fig. 1a, Supplementary Table 1). Alongside the involvement of canonical fibrotic pathways, we observed a significant enrichment of metabolic pathways, predominantly glucose-related, and of post-translational protein modifications within this subpopulation. To further investigate the role of metabolic dysregulation in IPF, we utilised a 3D organoid system, derived from lung tissue samples obtained from IPF patients and healthy, control resections from tumour explants. We first isolated EPCAM+ distal airway epithelial cells through tissue dissection and multiple sorting steps to obtain a pure epithelial cell fraction (Fig. 1b). The harvested cells were cultured in Cultrex basement membrane extract to promote the formation and growth of distal airway organoids (AOs)^15^. Under these culture conditions, AOs displayed a uniform, spherical morphology with a lumen and exhibit a pseudostratified epithelial layer encompassing functional basal, club, goblet, and ciliated cells (Fig. 1c, Extended Fig. S1a-c). Markers for basal (KRT5+), goblet (MUC5B+), and club cells (SCGB1A1+) were detected within 14 days of development (Fig. 1c, Extended Fig. S1 a), while cilia (FOXJ1+) became visible after 21 days (Extended Fig. S1a, b). Intriguingly, a reduction in ciliated area was observed in IPF AOs compared to control (Fig. 1d).

**Figure 1:**
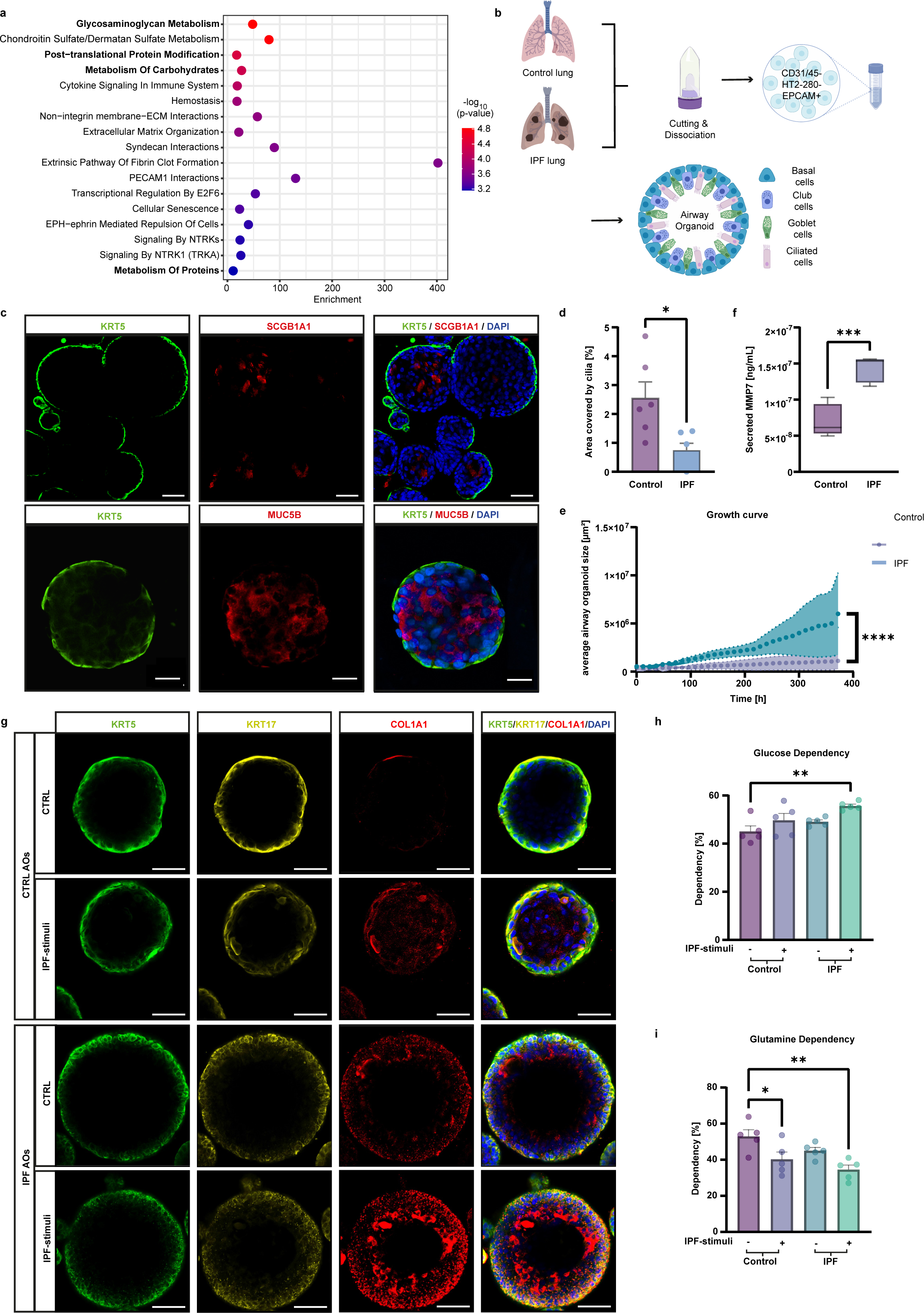
IPF-derived distal airway organoids exhibit aberrant basal cell characteristics. **a,** Reactome analysis of gene signatures in aberrant basal cells shows a strong inclination towards glucose-related metabolic pathways. **b**, Schematic representation of the cell isolation process from human distal lung tissue to generate patient-derived AOs. **c**, Representative immunofluorescence staining of 15 days AOs shows a pseudostratified epithelium with basal cells (KRT5+, green) externally, and differentiated club cells (SCGB1A1+, red) and goblet cells (MUC5B+, red) internally. Nuclei are stained with DAPI (blue). Scale bars are 20 µm (first row) and 50 µm (second row). **d**, Quantification of ciliated cell area indicates a significant decrease in cilia in IPF AOs from n = 6 IPF and n = 6 control AO donors (mean + s.e.m, \**p* < 0.05, unpaired *t*-test). **e**, Organoid size quantification over 15 days shows an increased proliferative capacity in IPF AOs (n = 4 IPF and n = 7 control AO donors, mean + s.e.m, \*\*\*\**p* < 0.0001, unpaired *t*-test). **f**, Profibrotic biomarker MMP7 secretion is increased in IPF AOs (n = 5 IPF and n = 5 control AO donors, mean + s.e.m., \*\*\**p* < 0.001, unpaired *t*-test). **g**, Representative immunofluorescence staining of AOs reveals aberrant basal cells (KRT5+(green)/KRT17+ (yellow)/COL1A1+ (red)). Nuclei are stained with DAPI (blue). Scale bars: 50 µm. IPF-stimuli further enriches this population after 7 days. **h, i**, Seahorse XF Mito Fuel Flex Test kit showed increased glucose (**h**) and decreased glutamine (**i**) dependency of IPF and IPF-stimulated AOs (n = 5 IPF and n = 5 control donors, mean + s.e.m., \**p* < 0.05, \*\**p* < 0.01, ANOVA/ Tukey’s).

Through time-lapse imaging, we noted that IPF AOs exhibited rapid initial growth (Fig. 1e, Extended Fig. S1d, e), reflecting the progressive nature of fibrotic disease^10,16^. Additionally, nanoindentation measurements on cells derived from IPF AOs showed increased stiffness of the epithelium (Extended Fig. S1f), falling within the range of an IPF lung^17,18^. This was accompanied by elevated levels of secreted MMP7, an IPF clinical biomarker^19^ (Fig. 1f).

Importantly, aberrant basal cells were detected in IPF AOs through immunofluorescence staining for KRT5, KRT17 and COL1A1, in contrast to control AOs (Fig. 1g). This phenotype was intensified upon exposure to pro-fibrotic stimuli^20^. Moreover, we identified a metabolic transition favouring heightened glucose dependency and diminished glutamine dependency (Fig. 1h, i, Extended Fig. S1g), as expected in fibrotic disease^21–23^.

In summary, our findings demonstrate the successful generation of functional patient-derived organoids that closely reflect key hallmarks of IPF, including aberrant basal cells, metabolic dysregulation, disease-associated biomarkers, and elevated matrix production, providing a valuable platform for studying disease mechanisms.

### IPF progression is associated with increased O-GlcNAcylation in aberrant basal cells

To delve deeper into the molecular mechanisms governing aberrant basal cell fate, we analysed transcriptional profiles of IPF and control AOs using RNA sequencing. Upon evaluation of the transcriptomic profile, we observed distinct clustering of the respective conditions and identified many IPF-associated differentially expressed genes (DEGs; p*-*value < 0.05) in IPF AOs (Fig. 2a, b, Supplementary Table 2a). This divergence was clearly enhanced upon exposure to pro-fibrotic stimuli (Extended Fig. S2a-c). Using an unbiased phenotypical signatures analysis, we observed an overrepresented signature of secretory cells characterised by SCGB1A1+/SCGB3A2+/SCGB3A1+^24^ in IPF AOs (Fig. 2c, Supplementary Table 2b). Notably, these cells have been previously described to be prevalent as a transitional state within severely fibrotic regions of the lung^25–27^. This prompts us to investigate the resemblance of the IPF AOs to human diseases by incorporating clinical data from various stages of IPF^28^. Gene set enrichment analysis (GSEA) unveiled a significant enrichment of gene signatures associated with IPF transitional alveolar type 2 (AT2)^14^ and lung profiles from patients with moderate IPF^28^ (Fig. 2d, Supplementary Table 2c). Together, this data corroborates the relevance of the IPF AO model to IPF, particularly in term of the classic histological description of the bronchiolization process.

**Figure 2:**
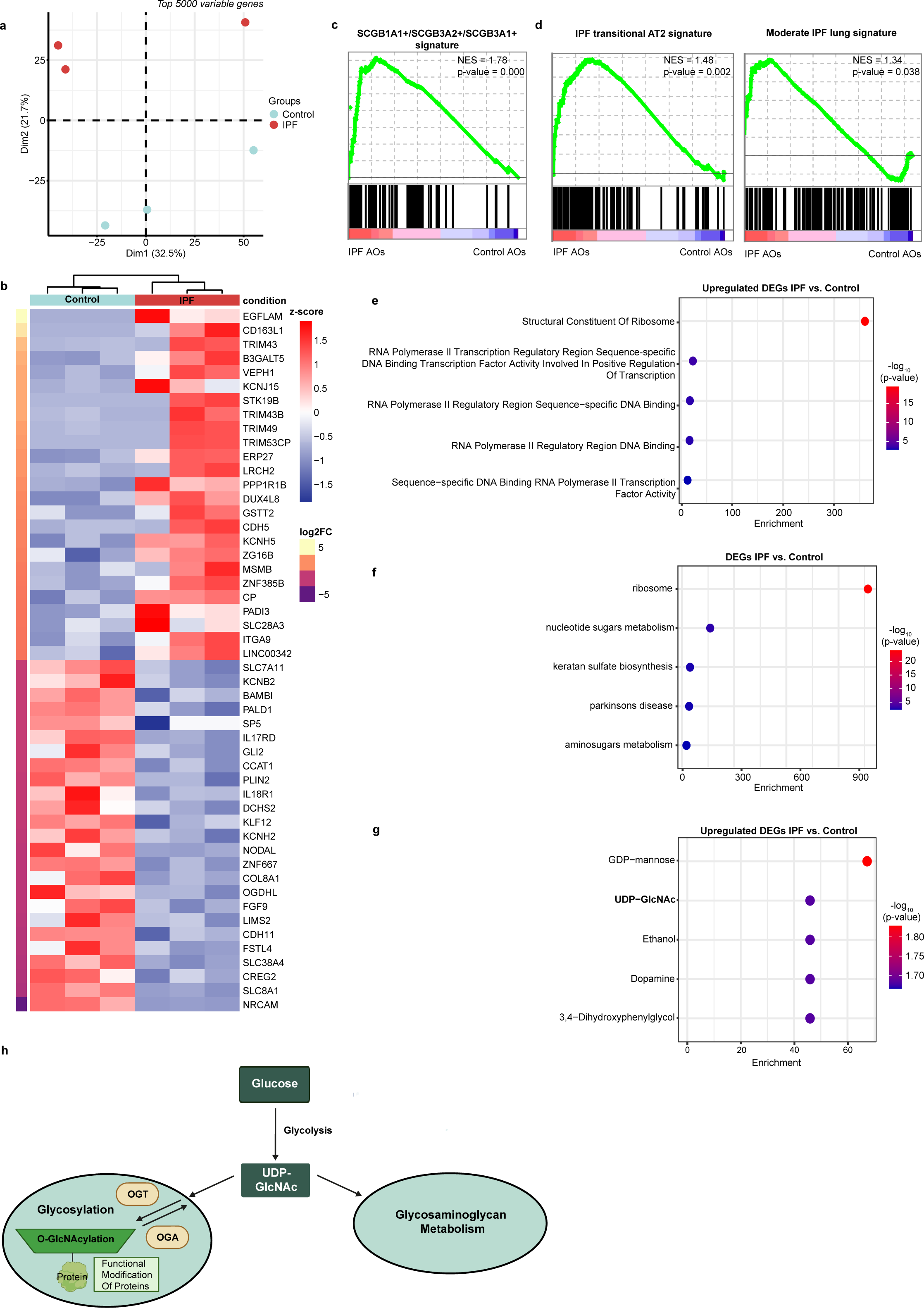
O-GlcNAc plays an important role in aberrant basal cells in IPF. **a**, Principal component analysis (PCA) of RNA-seq data with the top 5000 variable transcripts shows distinct clustering of IPF and control AOs. **b**, Hierarchical clustering of the top 50 differentially expressed genes (DEGs) from RNA-seq between IPF and control AOs (p-value < 0.05). **c**, GSEA using cell type signature database (c8) shows enrichment of a secretory signature (SCGB1A1+/SCGB3A2+/SCGB3A1+) in IPF AOs (p-value < 0.05). **d**, GSEA shows enrichment of an IPF transitional AT2 signature and moderate IPF lung stage signature in IPF AOs using IPF database (p-value < 0.05). **e**, Gene ontology (GO) molecular function analysis shows enrichment of increased transcriptional machinery activity in IPF AOs (p-value < 0.05, log2FC > 0). **f**, KEGG analysis shows enrichment of metabolic pathways, specifically glycosylation-related, among all DEGs in IPF AOs (p-value < 0.05). **g**, Metabolomics workbench metabolites analysis shows enrichment of UDP-GlcNAc metabolite in IPF AOs (p-value < 0.05, log2FC > 0). **h**, Schematic overview of UDP-GlcNAc and its relation to the glycosaminoglycan metabolism and glycosylation comprising O-GlcNAcylation. The precursor UDP-GlcNAc is reversibly added to proteins via OGT and OGA to induce O-GlcNAcylation on targeted proteins.

Next, we proceeded to explore the differences in gene expression between the IPF and control AOs. Among the most significantly enriched gene ontology (GO) terms in the upregulated DEGs of IPF AOs were those related to RNA polymerase II transcription machinery and transcription factors activities (Fig. 2e, Supplementary Table 2d). This indicates a potential transcriptional reconfiguration in IPF. Consistent with the scRNA-seq findings from IPF lungs (Fig. 1a), pathway analysis further revealed the engagement of various metabolic pathways in IPF AOs, particularly those associated with glycosylation and glycosaminoglycan metabolism (Fig 2f, Extended Fig. S2d, e and Supplementary Table 2e). This is intriguing as these metabolic processes are known to mediate transcriptional reprogramming through several routes, including the post-translational modifications (PTMs) of transcription factors^29^. This prompted us to delineate the specific PTMs induced by these metabolic pathways. Through metabolic regulation analysis that clusters gene sets by their common metabolites, we identified an enrichment of UDP-GlcNAc in IPF AOs (Fig. 2g, Supplementary Table 2f). This metabolite is a pivotal precursor for O-GlcNAcylation; a unique and reversible type of glycosylation PTM, and a building block in glycosaminoglycan metabolism processes^30,31^ (Fig. 2h). Collectively, these findings propose that dysregulation of O-GlcNAc modifications would modulate the aberrant cell behaviour, thereby exacerbate the progression of IPF specifically.

### Increased O-GlcNAc levels drive IPF progression

To test this hypothesis, we measured global O-GlcNAc levels in patient-derived lung lysates and detected a remarkable increase in IPF lungs (Fig. 3a). To access the regulatory role of increased O-GlcNAc levels in IPF, we deleted OGT (Extended Fig. S3a). As expected, OGT depletion attenuated the expression of pro-fibrotic genes in IPF-stimulated airway epithelial cells (Fig. 3b, c). Similar effects were observed within IPF-induced fibroblasts (Extended Fig. S3b, c, d), indicating a conserved regulatory role of aberrant O-GlcNAc levels in IPF cell fate decisions.

**Figure 3:**
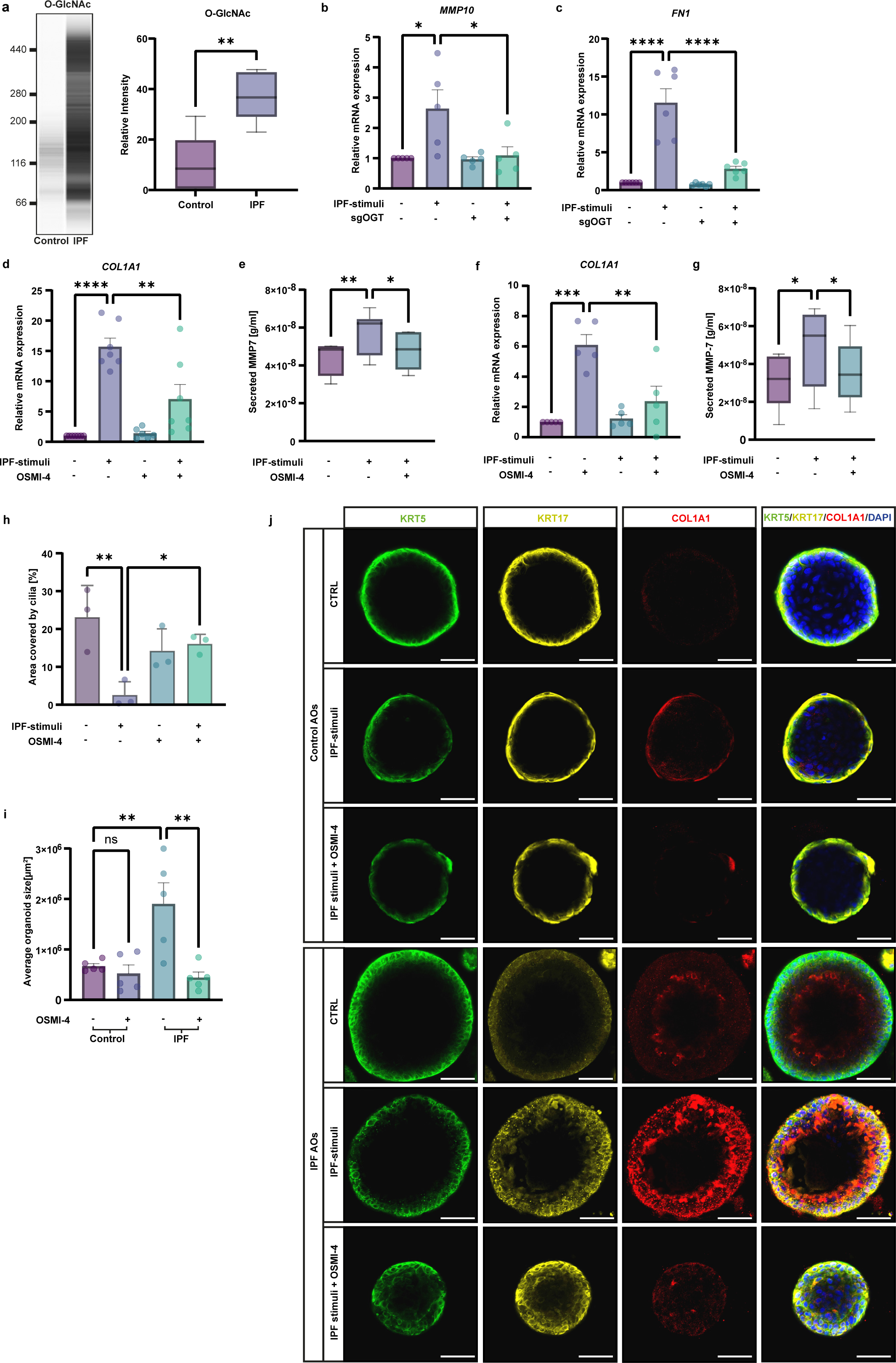
OGT inhibition reduces fibrotic characteristics and aberrant basal cell fate. **a**, Representative western blot analysis and quantification (right panel) of total O-GlcNAc levels in control (n = 6) and IPF (n = 5) lung lysates revealed significant increase of O-GlcNAc in IPF lungs (violin plot, \*\**p* < 0.01, unpaired *t*-test). **b**, **c**, RT-PCR analysis of *MMP10* (**b**) and *FN1* (**c**) in OGT-deleted airway epithelial cells treated with IPF-stimuli shows that OGT is required for profibrotic genes expression (n = 6, mean + s.e.m, \**p* < 0.05, \*\*\*\**p* < 0.0001, ANOVA/Tukey’s). **d,** RT-PCR analysis of *COL1A1* in IPF AOs treated with IPF-stimuli and OSMI-4 for 7 days shows decrease of gene expression upon OGT inhibition (n = 7, mean + s.e.m, \*\**p* < 0.01, \*\*\*\**p* < 0.0001, ANOVA/Tukey’s). **e**, ELISA analysis shows a decline in MMP7 secretion in an OGT-dependent manner upon stimulation with IPF-stimuli (n = 8, mean + s.e.m, \**p* < 0.05, \*\**p* < 0.01, ANOVA/Friedman). **f**, RT-PCR analysis shows that OGT inhibition attenuates chronically injured epithelial-fibroblast coculture induced *COL1A1* expression levels (n = 5, mean + s.e.m, \*\**p* < 0.01, \*\*\**p* < 0.001, ANOVA/Tukey’s). **g**, ELISA analysis reveals that OGT inhibition in chronically injured epithelial-fibroblast coculture ameliorates MMP7 secretion (n = 5, mean + s.e.m, \**p* < 0.05, ANOVA/Tukey’s). **h,** Quantification of ciliated cell area upon OSMI-4 treatment and chronic injury in epithelial-mesenchymal coculture increased area covered by ciliated cells (n = 3, mean + s.e.m, \**p* < 0.05, \*\**p* < 0.01, ANOVA/Tukey’s). **i**, Quantification of average organoid size shows decreased organoid growth upon inhibition of OGT after 10 days (n = 5, mean + s.e.m, \*\**p* < 0.01, ANOVA/Tukey’s). **j**, Representative immunofluorescence staining of IPF and control AOs treated with IPF-stimuli and OSMI-4 for 7 days shows decrease in aberrant basal cell signature upon OGT inhibition (scale bar 50 µm).

Subsequent to our hypothesis, we sought to explore the therapeutic potential of modulating OGT activity by employing the small molecule inhibitor OSMI-4. First, we tested the effect of this inhibitor on O-GlcNAcylation levels in 2D and 3D distal airway epithelial cultures and determined half-maximal effective concentrations (EC50) of 3 µM and 10 µM, respectively (Extended Fig. S3e-h). Intriguingly, inhibition of O-GlcNAc modifications by OSMI-4 culminated in a marked diminution of pro-fibrotic marker expression and secretion in both, IPF AOs (Fig. 3d, e) and epithelial-fibroblasts co-cultures (Fig. 3f, g, Extended Fig. S3i). Furthermore, OSMI-4 treatment effectively mitigated and restored the loss of ciliated cells in IPF (Fig. 3h). Notably, the proliferative capacity of IPF AOs was significantly attenuated upon OSMI-4 treatment (Fig. 3i). This observation aligns with previous findings which show anti-proliferative effects of inhibited O-GlcNAc levels in a variety of cancer cells^32,33^.

The incidence of aberrant basal cells was attenuated upon OSMI-4 treatment in both IPF and IPF stimuli-induced AOs (Fig. 3j). Remarkably, treatment with OGT inhibition in the presence of an IPF stimuli in IPF AOs led to a substantial reduction in aberrant basal cells, achieving levels comparable to those observed in control organoids.

Collectively, these experiments suggest the central role of O-GlcNAcylation in governing the aberrant cell fate decisions in IPF. The pharmacological inhibition of this post-translational modification by OSMI-4 not only mitigates fibrotic features but also appears to restore the epithelial integrity by reducing metaplastic aberrant basal cells and rescuing the loss of ciliated cell populations, thereby providing a compelling therapeutic avenue for ameliorating the pathological manifestations of IPF.

### IPF is associated with increased chromatin accessibility at promoter regions of metabolic genes

To delineate the fundamental alterations within the chromatin landscape that contribute to the O-GlcNAc-induced aberrant cell fates in IPF, we profiled the chromatin accessibility profiles using Assay for Transposase-Accessible Chromatin sequencing (ATAC-seq) in IPF and control AOs on a single-cell level focusing on the basal cell fraction (Extended Fig. 4a-c). Following the read alignment process and subsequent peak calling, we assessed the distribution of peaks with respect to established genomic features (Fig. 4a, b, Supplementary Table 3a). Despite the similarity in peak distribution across different conditions, the proximal promoter and transcription start site (within 1 kb) regions exhibited significantly greater accessibility in IPF AOs compared to control (48.89 % and 27.74 %, respectively).

**Figure 4:**
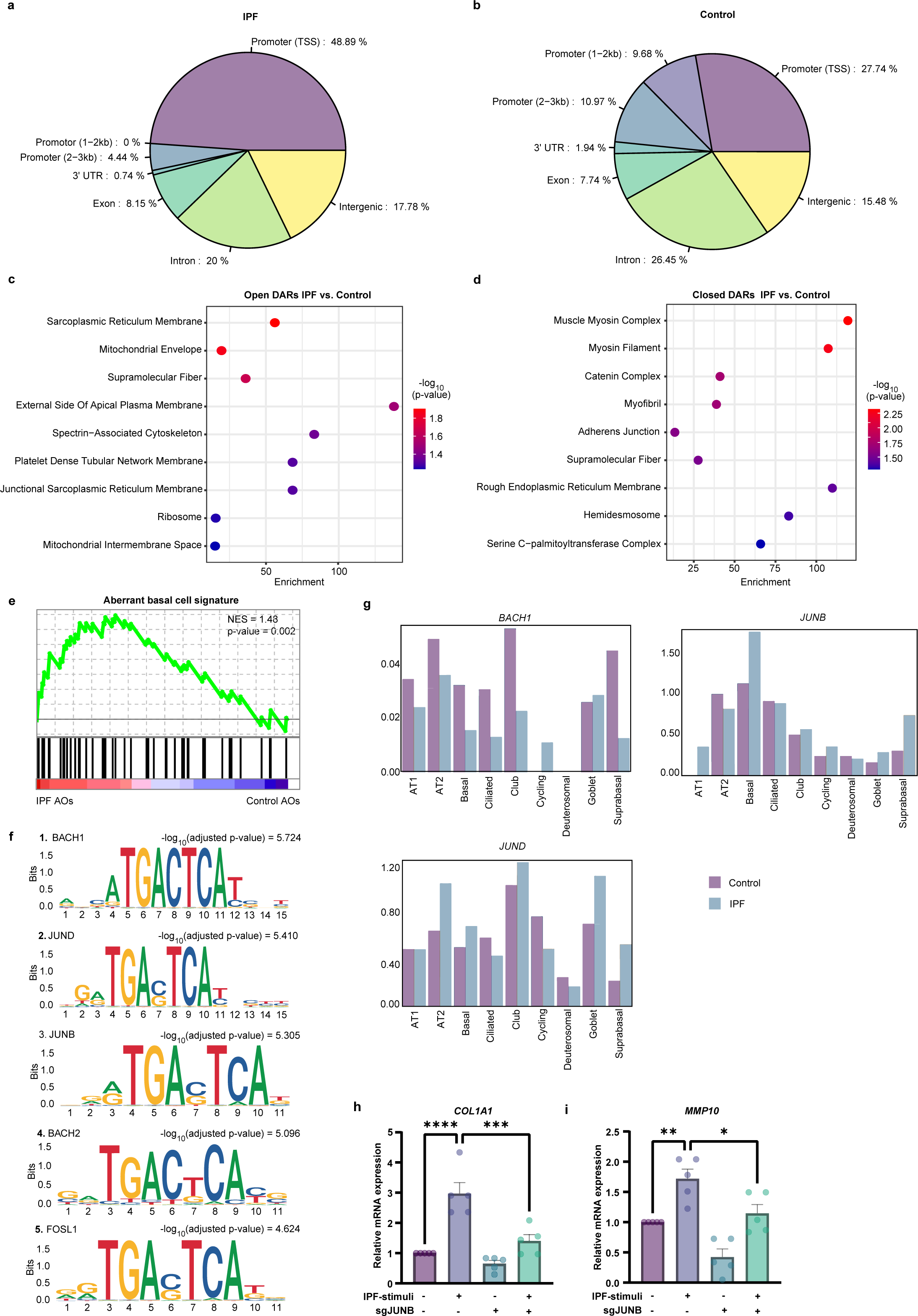
JUNB is enriched at promoter regions of open chromatin in IPF. **a**, **b**, Genomic annotations of IPF basal cells in AOs for significantly differentially open (**a**) and closed (**b**) accessible regions (DARs, p*-value* < 0.05) shows enrichment of open chromatin in IPF at promotor and transcription start site (TSS) regions. Genomic regions are defined as introns, exons, intergenic regions, 3’ UTR, promotor TSS (defined as peak summits within 1 kb upstream and downstream of the TSS), promotor (1-2 kb), and promotor (2-3 kb). **c**, **d**, Gene ontology (GO) enrichment analysis of open (**c**) and closed (**d**) DARs using annotated genes (*q*-value < 0.05) in IPF basal cells. Open regions were associated with signal transduction and metabolic pathways, whilst closed regions were related to mechanical stress and barrier function. **e**, GSEA reveals enrichment of annotated genes based on DARs (*q-value* < 0.05) for aberrant basal cell signature in IPF basal cells. **f**, Top 5 transcription factor (TF) motif enrichments in IPF basal cells. **g**, Expression levels of the top TFs in different epithelial cells. Note, *JUNB* displays highest differential expression in IPF basal cells. **h, i**, RT-PCR analysis shows attenuated the expression of profibrotic genes after fibrotic induction with IPF-stimuli in airway epithelial cells (n = 5, mean + s.e.m, \**p* < 0.05, \*\**p* < 0.01, \*\*\**p* < 0.001, \*\*\*\**p* < 0.0001, ANOVA/Tukey’s).

GO enrichment analysis of significantly differentially accessible regions (DARs) revealed that IPF AOs were associated with decreased chromatin accessibility at promoter regions of genes responsible for mechanical stretch and barrier function (Fig. 4d, Supplementary Table 3b), whereas regions associated with metabolism and signal transduction gained accessibility (Fig. 4c, Supplementary Table 3c). Additionally, these altered accessible regions showed a significant correlation with the IPF aberrant basal cell signature (Fig. 4e, Supplementary Table 3d), suggesting a reprogramming of gene regulatory network that may offer insights into disease etiology.

Next, to gain insight into the regulatory network of open chromatin accessibility in IPF aberrant basal cells, we conducted transcription factor (TF) motif enrichment analysis and found significantly overrepresented motifs specific to the activator protein 1 complex in the IPF condition (Fig. 4f). The presence of these TFs, which have been reported to play essential roles in regulating gene expression in response to stress^34–36^, implicates their contribution to the pathophysiology of IPF. Similar findings were also observed in a larger cohort of IPF donors within using a bulk ATAC-seq analysis (Extended Fig. S4d-k, Supplementary Table 3e).

We further assessed whether the expression of these TF motifs exhibited differential expression in IPF and found that JUNB was significantly upregulated in IPF aberrant basal cells^37^ (Fig. 4g). Remarkably, the depletion of JUNB attenuated the pro-fibrotic gene expression in both, IPF-treated airway epithelial cells and fibroblasts (Fig. 4h, i and Extended Fig. S4l, m). Collectively, our findings revealed that IPF is characterised by increased chromatin accessibility and gene expression, particularly at promoter regions with the JUNB motif that control pro-fibrotic cell fate decisions.

### O-GlcNAcylation determines pro-fibrotic cell fate decisions through JUNB

Previous studies discovered that TFs could carry O-GlcNAc modifications, which influence their transcriptional activity and stability^29^, therefore we hypothesise that O-GlcNAcylation of JUNB may serve as a stress sensor to regulate pro-fibrotic response in the lung. To evaluate this hypothesis, we examined the presence of O-GlcNAc modification on JUNB and its role in IPF. As expected, immunoprecipitation (IP) showed an increase in O-GlcNAc PTM levels on JUNB upon pro-fibrotic stimulation in airway epithelial cells and fibroblasts (Fig 5a, Extended Fig. S5a). To validate the specific role of O-GlcNAcylated JUNB in the pro-fibrotic response, we substituted the potential O-GlcNAc sites on JUNB^38^ from serine to alanine (S-to-A) and threonine to valine (T-to-V) via site-specific mutations^39^, and expressed this loss-of-function (LOF) of O-GlcNAc sites on JUNB in airway epithelial cells (plasmid sequence in Supplementary Table 4). This led to an attenuation in the expression of pro-fibrotic genes and biomarker MMP7 upon IPF-stimuli (Fig. 5b-d, Extended Fig. S5b-d). Furthermore, expression of JUNB-LOF in human primary fibroblasts treated with IPF-stimuli exerts a reduction in the pro-fibrotic response, suggesting the effect of O-GlcNAcylated JUNB extends beyond epithelial cells (Extended Fig. S5e-g).

**Figure 5:**
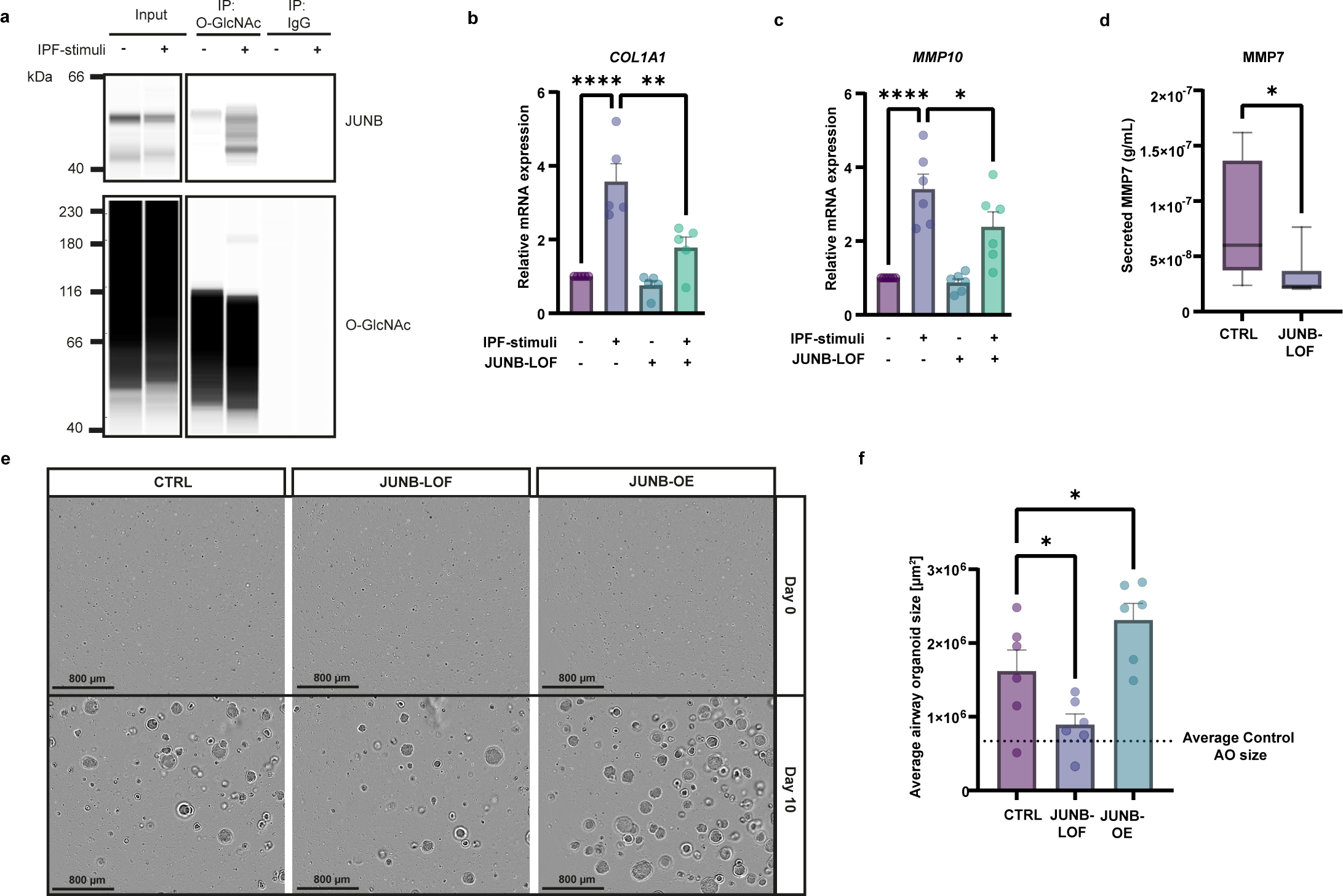
O-GlcNAcylation of JUNB induces pro-fibrotic cell fate decisions. **a**, Representative western blot analysis of the O-GlcNAc immunoprecipitated fraction shows increased levels of O-GlcNAc mark on JUNB in injured airway epithelial cells (n = 3 independent donors). IgG antibody was employed as negative control. **b**, **c**, RT-PCR analysis shows diminished expression of pro-fibrotic genes (*COL1A1* (**b**) and *MMP10* (**c**)) upon transfection of JUNB-LOF (loss-of-function) and fibrotic induction IPF-stimuli in airway epithelial cells (n = 5, mean + s.e.m, \**p* < 0.05, \*\**p* < 0.01, \*\*\*\**p* < 0.0001, ANOVA/Tukey’s) Plasmid was generated by site-specific mutations in the O-GlcNAc sites of JUNB. **d**, Transduction of JUNB-LOF plasmid in n = 6 IPF AOs led to a decrease in the secretion of MMP7 after 10 days (mean + s.e.m, \**p* < 0.05, Wilcoxon). **e**, **f**, Representative pictures (**e**) and quantification (**f**) of average organoids size of n = 6 IPF AOs was reduced after JUNB-LOF transduction in AOs after 10 days, whereas overexpression of JUNB led to a significant increase in growth (mean + s.e.m, \**p* < 0.05, ANOVA/Tukey’s).

We next assessed the role of O-GlcNAcylated JUNB using IPF AOs. We evaluated the average organoid growth over a 10-day period post adeno-associated viral (AAV) transduction with a JUNB-LOF construct. Additionally, we also incorporated an overexpression of JUNB (JUNB-OE). Notably, JUNB-LOF resulted in a significant reduction in organoid size after 10 days reaching levels similar to those seen in control AOs, while overexpression stimulated organoid growth in IPF AOs (Fig. 5e, f). Together, these data underline the involvement of O-GlcNAcylated JUNB in IPF pathology, particularly its pivotal role in steering the pro-fibrotic response in the lung.

### Increased O-GlcNAcylation levels compromise alveolar differentiation *ex vivo*

Finally, we investigated the impact of this machinery rheostat within the context of more complex native lung tissue. Therefore, we employed an *ex vivo* model of rat precision cut lung slices (rPCLS). This system retains the intricate cellular diversity and spatial architecture inherent to the lung, thereby providing a robust platform for pharmacological investigations^40^. We induced fibrosis in healthy rat lung tissues through the administration of a rat-specific IPF-relevant cocktail (r-IPF-RC)^41^.

Following a 24-h incubation period post slicing, rPCLS were subjected to the respective treatment with r-IPF-RC and OSMI-4 for a total of 120 h (Fig. 6a). We first verified that the elevated O-GlcNAc levels in fibrosis were preserved in rat lung tissues (Extended Fig. S6a). Importantly, the induction of a fibrotic phenotype was successfully achieved as evidenced by an upregulation in pro-fibrotic and mucus-associated gene expression (Fig. 6b-d) and a concurrent decrease in the AT2 cell marker surfactant protein c (*Sftpc)* (Fig. 6e).

**Figure 6:**
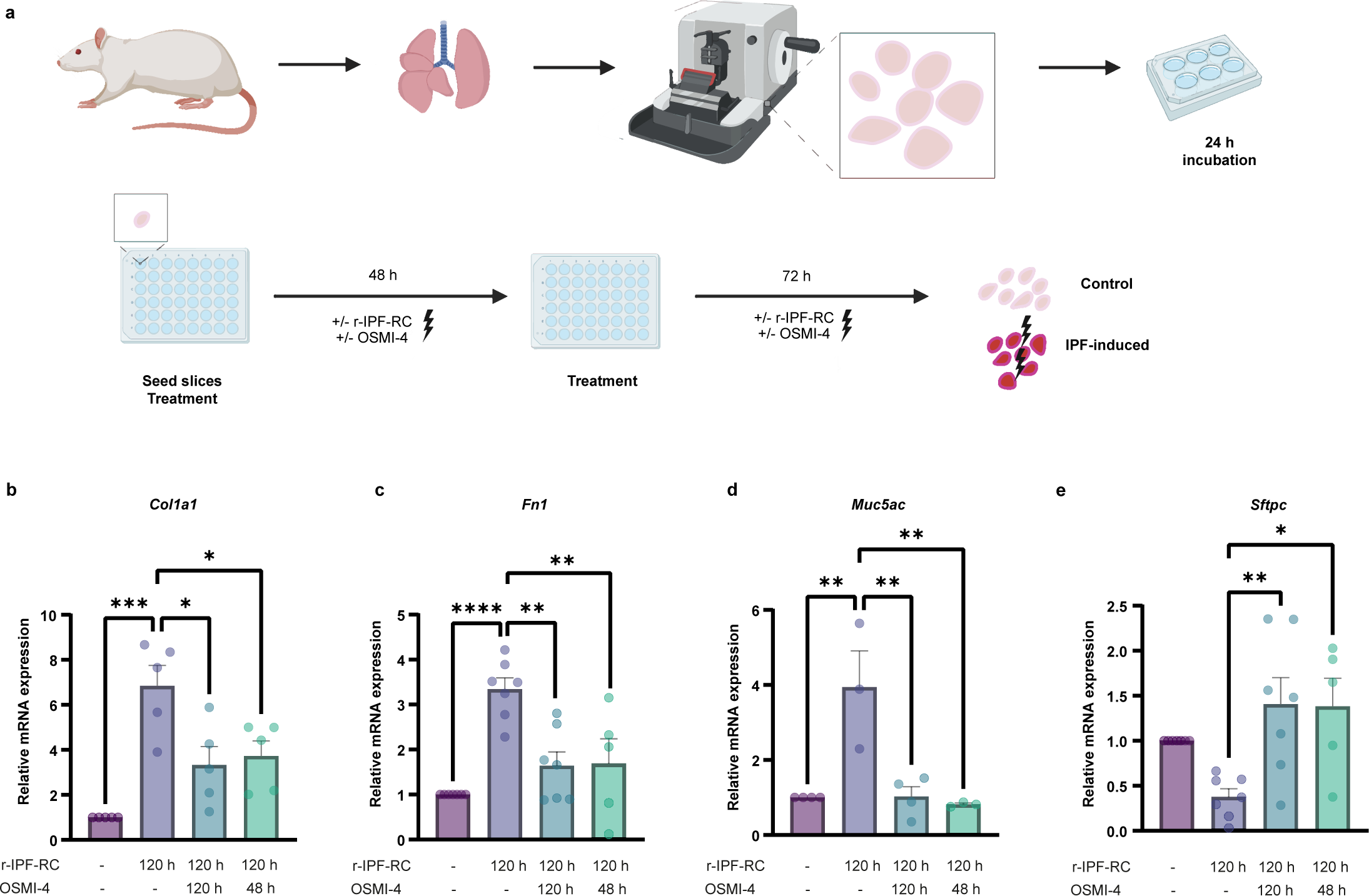
O-GlcNAcylation mediates epithelial cell fates in an *ex vivo* model. **a**, Schematic overview of generation and treatment of rPCLS. Healthy rats were euthanized, lungs were filled with agarose and lung tissue dissected. Slices were cut with a microtome to a thickness of 200-350 µm. Afterwards, slices were incubated for 24 h prior to seeding and treatment with +/- r-IPF-RC and +/- OSMI-4 for 48 h followed by additional treatment for another 72 h (total treatment time: 120 h). **b** -**e**, RT-PCR analysis of *Col1a1* (**b**), *Fn1* (**c**), *Muc5ac* (**d**), and *Sftpc* (**e**) in rPCLS with r-IPF-RC and with OSMI-4 for 120 h showed reduction in fibrotic response and induction of regeneration compared to control (n = 5 rats, 2 slices per rat, mean + s.e.m, *p < 0.05, **p < 0.01, ***p < 0.001, ****p < 0.0001, ANOVA/Tukey’s).

Simultaneous inhibition of O-GlcNAc for 120 h mitigated the pro-fibrotic and mucous response and augmented *Sftpc* expression (Fig. 6b-e), suggesting the prevention of IPF hallmarks by OSMI-4. Remarkably, this effect was also replicated in a therapeutic setting with OSMI-4 added only in the last 48 h of the experiment. This finding suggests that reducing O-GlcNAc levels in pulmonary fibrosis not only diminishes pro-fibrotic factors, but also enables alveolar repair and regeneration.

## Discussion

In the disease state, cells adopt a unique metabolic state, distinct from their healthy counterparts, to sustain their survival in a hostile pathologic microenvironment^42^. This adaptation involves the use of various metabolic pathways across different cell types, indicating a complex regulatory mechanism driving disease progression. In this study, we demonstrate how a metabolic shift impacts chromatin accessibility and transcriptional reprogramming of aberrant basal cells in driving the bronchiolization process in IPF, as summarised in figure 7.

**Figure 7:**
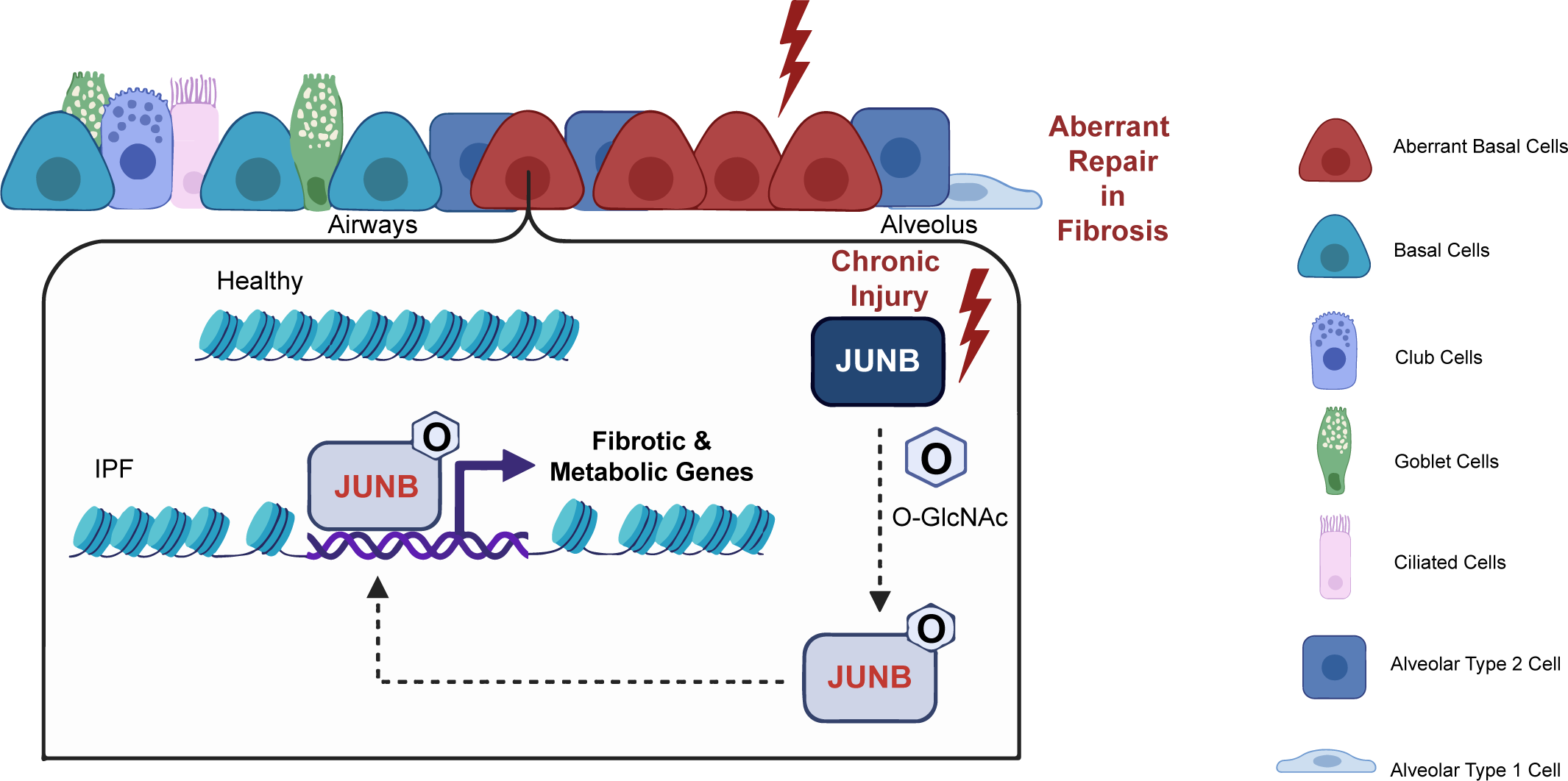
Schematic model describes the expansion of aberrant basal cells driven by the O-GlcNAC-JUNB axis. Chronic injury within the lung epithelium fosters an aberrant basal cell phenotype, which in turn elevates global O-GlcNAc levels, and O-GlcNAcylation of JUNB. The O-GlcNAc-JUNB drives the fibrotic cell fate by accessing open promotor regions of metabolic and pro-fibrotic genes to promote aberrant repair and fibrosis.

While human organoids have commonly been employed to study genetic diseases, our data highlights the use of a 3D *in vitro* organotypic disease model for IPF, derived from patient tissue with non-genetic upstream pathological stimuli. Remarkably, this model successfully recapitulates important hallmarks of IPF, including increased tissue stiffness, the emergence of aberrant basal cells and metabolic dysregulation^43,44^. Interestingly, tissue mechanics are known to influence the chromatin landscape and thus stem cell function in ageing^45^ and age-related diseases^46^. To the best of our knowledge, chromatin landscape profiling for epithelial bronchiolization in IPF has not been analysed. Our findings suggest that tissue stiffening could trigger a series of molecular events, including a metabolic shift crucial for stem cell functions such as activation, proliferation, and migration. This is regulated through the O-GlcNAc-JUNB axis, which enhances chromatin accessibility and transcriptional reprogramming, ultimately leading to the aberrant behaviour and metaplastic proliferation of KRT5+ basal cells in alveolar regions.

Elevated O-GlcNAc levels are a hallmark of a variety of diseases^30^, including fibrosis^47^ and cancer^48^, yet their precise targets are elusive. Here, we observe increased O-GlcNAc levels in the bronchiolization process in IPF. Together with previous research on fibroblasts^47^, our findings indicate that O-GlcNAcylation may play a crucial role in the development and progression of fibrosis throughout the lung. Importantly, JUNB, a part of the activator protein 1 transcription factor family, has been implicated in a myriad of biological processes^49–52^ and disease pathogenesis, particularly in cancer^35,53^. However, its role is controversial, and has been classified both as a tumour suppressor^54^ and an oncogene^35,36^. Our findings introduce a concept that O-GlcNAcylation may fundamentally alter cellular activity of JUNB, shedding new light on the connection between metabolism and stress response. Depending on its O-GlcNAc status, it may switch JUNB activity across diverse environmental conditions, as it was previously observed for MORC2^55^. Intriguingly, inhibiting the O-GlcNAc modification on JUNB attenuates the metaplastic differentiation of KRT5+ aberrant basal cells and induces regenerative features, such as the decrease of aberrant basal cells in IPF AOs and the re-emergence of alveolar cell markers in rPCLS. The therapeutic benefits seen from using OSMI-4 to block global O-GlcNAc levels could extend beyond the effects on JUNB; the role of O-GlcNAc-modifications in the context of IPF is still largely unexplored and warrants further investigations. This highlights the importance of future studies aimed at mapping the O-GlcNAcome implicated in IPF. Additionally, the development of more targeted inhibitors for O-GlcNAcylation of JUNB could pave the way for novel treatment strategies.

Collectively, our study sheds light on how metabolic changes might convert into established signalling pathways through O-GlcNAc, which serves as a modulator of JUNB, promoting fibrotic cell fate decisions and aberrant basal cell invasion. This insight offers a new avenue for selectively targeting this specific modification, as opposed to broadly inhibiting TF activity, presenting a viable therapeutic approach to arrest fibrotic progression and stimulate lung regeneration.

## MATERIALS AND METHODS

### Ethical approval

Patient lung tissue was obtained from the German Centre for Lung Research (DZL/BREATH) Hannover in Germany with full local ethical approval by DZL’s ethic committee and patient consent. Further airway organoid donors were purchased at HUB Organoids as ready-to-use cell stocks with ethical approval and patient consent.

All human tissue samples are coded / de-identified. Studies were performed in accordance with the local legislation and institutional requirements.

### Human lung tissue processing and AO cultivation

To isolate human lung cells, tissue samples are collected from the distal lung and cut into 2 mm slices. Subsequently, the tissues are enzymatically treated and placed into the gentleMACS dissociator to induce cell detachment. From the obtained cell suspension dead cells are removed and erythrocytes are depleted. Eventually, the remaining cells are magnetically sorted out with MACS. Firstly, endothelial cells (CD31/45+ cells) are collected. From the remaining cell suspension, alveolar cells (HT2-280+ cells) are sorted out, yielding the last fraction from which the distal airway epithelial cells are obtained (EpCAM+ cells). Next, the EpCAM+ cells are seeded in Cultrex RGF BME type 2 with airway organoid media to support organoid formation and expansion (based on Sachs et al.^15^). Once organoids have fully formed, they are cultivated depending on the experimental set-up.

To further induce fibrosis in AOs, an IPF-relevant cytokine cocktail (IPF-RC)^20^ was used 5x concentrated for 7 days. Composition of 5x IPF-RC is listed in Supplementary Table 5. To block OGT in AOs a commercially available inhibitor (OSMI-4) was used in 10 µM concentration for 7 days.

### 2D cultivation of primary cells

Healthy human lung airway epithelial cells (SAECs, Lonza, #CC-2547, or patient-derived distal airway epithelial cells) were used in passages 2-3 and expanded on type I collagen (Corning, #354236) pre-coated cell culture flasks in PneumaCult™-Ex Plus Medium (STEMCELL Technologies, #05040). For air-liquid-interface differentiation (ALI), airway epithelial cells were seeded on collagen-coated transwells, grown until confluency and airlifted by changing medium to PneumaCult™-ALI Medium (STEMCELL Technologies, #05001) and exposing cells apically to air. Differentiated cells were observed after 10 days.

Healthy human lung fibroblasts (LF) (Lonza, #CC-2512) were used in passages 4-8 and expanded in flasks in Fibroblast Basal Medium (Lonza, #CC-3131) with FGM2-Fibroblast Growth Medium BulletKit (Lonza, CC-3132).

For pro-fibrotic induction 3 ng/ml TGF-β1 and for OGT inhibition 3 µM OSMI-4 was used for 72 h.

### Epithelial-mesenchymal chronic injury coculture

SAECs were differentiated in air liquid interface (ALI) on transwells for 10 days. Afterwards, ALI SAECs were treated for 14 days with 1 ng/ml TGF-β1 or vehicle to induce chronic injury and cultivated in coculture with human lung fibroblasts +/- 1 ng/ml TGF-β1 in the basolateral chamber for additional 3 days.

### Histology

AOs were seeded and cultivated as previously described. After 25 days, AOs were dissolved away from Cultrex® RGM Type 2 (R&D systems, #3533-010-02) and fixed in 4 % paraformaldehyde (PFA) for 20 min at RT. Afterwards cells were washed and resuspended in HistoGel^TM^ (Richard-Allan Scientific, #HG-4000-012) and transferred to a paraffin cassette and incubated for 30 min at 4°C. Next, the cassette was embedded in paraffin and cut into 3 µm slices by using microtome. For subsequent Hematoxylin-Eosin or Periodic-Acid-Schiff staining, the respective AO sections were deparaffinised and further stained according to standard protocols.

### Immunofluorescence staining

For immunofluorescence staining, control and IPF AOs were seeded on µ-slide 8-well chamber slides (IBIDI, #80807), and cultivated or treated as previously described. AOs were fixed with 4 % PFA for 30 min and permeabilized with 0.2 % TritonX-100 in PBS, followed by blocking in 1 % BSA in PBS. Samples were incubated overnight at 4 °C with the respective primary antibody, washed with PBS and incubated with secondary antibodies and DAPI (Thermo Fisher, #62248). After final washing, samples were covered with PBS and imaged with either the LSM710 or LSM900 (Zeiss) confocal microscope using 20x, 40x, or 63x immersion objectives. Images were evaluated using ZEN software3.7 (Zeiss).

Primary antibodies used: mouse anti-Mucin-5AC CF488 conjugated (Invitrogen, #MA1-38223; 1:50) rabbit anti-Keratin-5 Alexa Fluor 647 conjugated (Abcam, #ab193895; 1:100); rat anti-SCGB1A1 (Novus Biological, #MAB4218-SP; 1:50); rabbit anti-KRT5 (Cell Signaling, #25807; 1:500); guinea pig anti-Keratin-17 (Progen, #GP-CK17, 1:300); mouse anti-Collagen 1a1 (Sigma Aldrich, #SAB4200678, 1:100).

Secondary antibodies used (all from Invitrogen): anti-rat immunoglobulin-G (IgG) Alexa Fluor (AF) 568 (#A-11077; 1:500); anti-rabbit IgG AF488 (#A-11034, 1:500); anti-guinea pig IgG AF568 (#A11075; 1:500); anti-mouse IgG AF647 (#A21235, 1:500).

### Nanoindentation

The stiffness of airway epithelial cells derived from AOs and seeded on transwells was measured via interferometry-based optical fibre-top nanoindentation by Pavone (Optics11 Life, Amsterdam, Netherlands). Cantilevers with stiffnesses of 0.029 N/m and 0.017 N/m and an area of 32.5 μm^2^ was applied. Prior to the measurement, a matrix scan was performed where nanoindentation of 30 points per well was performed.

### RNA extraction and quantitative RT-PCR

Total RNA was extracted using the RNeasy Plus Micro kit (Qiagen, #74034) and concentration measured by NanoDrop. Reverse transcription into cDNA was performed using the SuperScript™ VILO™ cDNA Synthesis Kit (Invitrogen, #11754050). Gene expression analysis was carried out by using TaqMan assays and TaqMan Advanced Master Mix (Applied Biosystems, #4444557). Samples were measured in triplicates on the Quantstudio 6 Real Time PCR machine. Changes in gene expression were determined using the comparative cycle threshold method with housekeeper genes RPS26, GAPDH, and B2M.

The following human (h) and rat (r) TaqMan assays (Thermo Fisher) were used: *hKRT5* (Hs00361185_m1); *hSCGB1A1* (Hs00171092_m1); *hMUC5B* (Hs00861595_m1); *hFOXJ1* (Hs00230964_m1); *hKRT17* (Hs00356958_m1); *hCOL1A1* (Hs00164004_m1); *hOGT* (Hs00269228_m1); *hMMP10* (Hs00233987_m1); *hFN1* (Hs01549976_m1); *hCOL5A1* (Hs00609133_m1); *hJUNB* (Hs00357891_s1); *hRPS26* (Hs00955682_g1); *hGAPDH* (Hs02786624_g1); *hB2M* (Hs00187842_m1); *rCol1a1* (Rn07363068_m1), *rFN1* (Rn00569575_m1); *rMuc5ac* (Rn01451252_m1); *rSftpc* (Rn00569225_m1); rGapdh (Rn01775763_g1)

### IncuCyte Organoid Growth Assay

To determine average organoid size over time, singularized organoids were seeded in a concentration of 1000 cells/µl and cultivated according to organoid cultivation protocol. For AAV6.2 induced JUNB-LOF, JUNB-OE, and GFP-control expression (Supplementary Table 5), AOs were seeded and subsequently transduced with AAV vectors for 24 h at a MOI of 10^5^.

Images were taken every 12 h for 10-15 days by using IncuCyte® Live Cell Analysis system with a 4x Plan-Apochromat objective in organoid assay mode. Analysis was conducted using IncuCyte® Live Cell Analysis software.

### ELISA

Supernatants were centrifuged and diluted 1:10. Measurement of human MMP7 (Biotechne, #DY907) was performed according to manufacturer’s instructions. Relative fluorescence intensity (excitation 490 nm, emission 520 nm) was measured via SpectraMax M5 and SoftMax ® Pro software (Molecular Devices). Total protein concentration was determined by standard curve.

### Cilia beating measurement

Active beating cilia were calculated from image stacks of 2D+time as previously described^56^. In brief, four different regions evenly distributed over the transwell were imaged using 32x objective of an Axiovert 25 microscope (Zeiss) and an acA 1300-200µm black and white USB-3.0 high speed camera (Basler). Movement of cilia was recorded by taking 100 frames per second for a total of 6 seconds per transwell area (total of 512 images with a size of 512*512 pixels). Applications for image and analysis were developed by using HALCON 13.0.2 (MVTec Software) machine vision software toolbox and to visualise the image stack Analyze (AnalyzeDirect) was applied. Area covered by active cilia was calculated by quantification of pixels which were detectably moving within the chosen area.

### Seahorse Mito Fuel Flex Test

Seahorse Mito Fuel Flux Assays were performed using XFe96 Bioanalyzer (Agilent). Airway epithelial cells were seeded on collagen-coated XFe96 cell culture plates and cultivated for 48 h. Afterwards, 5x IPF-RC was used for 72 h and the assay performed according to manufacturer’s instructions (Agilent, #103260-100). Calculations of Capacity [%] and Dependency [%] were performed using Wave Software V2.6.3.5 based on OCR measurements (Agilent).

### Western Blot

To determine protein expression levels, simple western automated western blot systems PeggySue, SallySue or Jess (Biotechne) were used according to manufacturer’s instructions (Protein simple, Biotechne, #SM-S001, #SM-S002). Quantification was performed by normalisation to calnexin or using Simple Western total protein quantification. To obtain proteins, tissue was lysed using T-Per tissue extraction buffer or cells using RIPA buffer with phosphatase/protease inhibitor, respectively. Antibodies used: mouse anti-OGlcNAc RL2 (Novus Biologicals, #NB300-534, 1:500), mouse anti-JUNB (Thermo Fisher, #SAB1406056, 1:50), rabbit anti-calnexin (Cell Signaling, #2679, 1:700).

### Immunoprecipitation

Cells were harvested in IP lysis buffer with phosphatase-/protease-inhibitor followed by short incubation and lysate clearance at 18,000 g. Protein concentration was determined using Pierce BCA assay (Thermo Fisher, # 23225). Pre-coupled antibody beads (Cell signaling, Custom Conjugation Service; rabbit anti-O-GlcNAc IgG (#82332) and rabbit IgG isotype control (#3900)) were washed twice and taken up in IP lysis buffer. Next, they were mixed with 500 µg protein per reaction and incubated at 4 °C overnight. After repetitive washing with IP lysis buffer, proteins were eluted using Simple Western 5 x Sample buffer and 5 x Master Mix (1:1) at 95 °C at 300 rpm for 5 min, followed by Simple Western analysis (PeggySue).

### RNA extraction and RNA-seq

After taking out the respective number of cells for ATAC-seq, cells were pelleted and lysed in RLT Plus buffer (QIAGEN, #1053393). Total RNA was extracted and purified on the MagMAX instrument (Thermo Fisher) using the MagMAX™ 96 Total RNA Isolation Kit (Thermo Fisher #AM1830) following manufacturer’s instructions. Total RNA was quantitatively and qualitatively assessed using High Sensitivity dsDNA Quanti-iT Assay Kit (ThermoFisher) on a Synergy HTX (BioTek) and High Sensitivity Total RNA Analysis DNF-472 on a 96-channel Fragment Analyzer (Agilent), respectively. All samples had RIN values >9. Total RNA samples were normalized on the MicroLab STAR automated liquid platform (Hamilton). A total RNA input of 250ng was used for library construction with the NEBNext Ultra II Directional RNA Library Prep Kit for Illumina (#E7760), together with the NEBNext Poly(A) mRNA Magnetic Isolation Module (#E7490) and NEBNext Multiplex Oligos for Illumina (#E7600) (New England Biolabs). RNA-seq libraries were processed on a Biomek i7 Hybrid instrument (Beckman Coulter) using 13 PCR cycles, following the manufacturer’s protocol, except for the use of Ampure XP beads (Beckman Coulter) for double-stranded cDNA purification and PCR clean-up. Final sequencing libraries were quantified by the High Sensitivity dsDNA Quanti-iT Assay Kit (ThermoFisher) on a Synergy HTX (BioTek). Libraries were also assessed for size distribution and adapter dimer presence (<0.5%) by the High Sensitivity NGS Fragment 1-6000bp kit on a 96-channel Fragment Analyzer (Agilent). All sequencing libraries were then normalized on the MicroLab STAR (Hamilton), pooled and sequenced on an Illumina NovaSeq 6000 using an S2 Reagent Kit v1.5 (Run parameters: 101bp, Rd2: 8bp, Rd3: 8bp, Rd4:101bp) with an average sequencing depth of ∼30 million pass-filter reads per sample.

### Analysis of bulk RNA-seq

Differential expression analysis was carried out with DESeq2 (v1.42.0)^57^. Statistical tests were performed via the Wald test and p-values were corrected with the Benjamini-Hochberg method. Genes with a p-value below 0.05 were considered differentially expressed (Supplementary Table 2a). Differential expressed genes were further analysed using EnrichR^58,59^ and GSEA^60^.

### Bulk ATAC sample preparation and sequencing

For matched bulk-ATAC and RNA-seq three IPF- and control-derived airway organoid donors were used, respectively, and singularized. ATAC-seq library prep was performed as previously described^61^ with minor adaptions. In brief, 70,000 cells of singularized AOs were lysed in lysis buffer (10 mM Tris pH 7.5, 10 mM NaCl, 3 mM MgCl2, 0.1 % NP-40) for 5 min and centrifuged. Pelleted cells were transposed using Tagment DNA TDE1 Enzyme and Buffer Kit (Illumina, # 20034197) for 60 min at 37 °C with shaking. Afterwards, samples were purified using clean-up buffer (0.5 M NaCl, 400 mM EDTA, 1 % SDS, 2 mg/ml Proteinase K) and normal phase 2x AMPureXP (Beckman Coulter, #A63881) bead clean-up, followed by PCR amplification. Finally, amplified samples were purified using DNA Clean & Concentrator Kit (Zymo, #ZYM-D4029) and double-sided clean-up with AMPure beads. Libraries were quantitatively and qualitatively assessed using the 1x dsDNA kit on the Qubit 4 Fluorometer (ThermoFisher) and the High Sensitivity NGS Fragment 1-6000bp kit on a 48-channel Fragment Analyzer (Agilent), respectively. ATAC-seq libraries were normalized on the MicroLab STAR (Hamilton), pooled and spiked in with PhiX Control v3 (Illumina). Libraries were sequenced on an Illumina NovaSeq 6000 using an S2 Reagent Kit v1.5 (#20028315; Read parameters: Rd1:101bp, Rd2: 8bp, Rd3: 8bp, Rd4:101bp) with an average sequencing depth of ∼100 million reads passed filter per sample.

### Analysis of bulkATAC-seq

After sequencing we used the nf-core ataq-seq pipeline (v2.1.2)^62,63^ to perform alignment and downstream analysis. Briefly, pre-alignment quality control and adapter trimming of the FASTQ files was performed with the default parameters. The reads were then aligned to the human reference genome (GRCh38.86) and alignments from multiple libraries per sample were merged. The standard settings of the pipeline were kept to remove low-quality reads, including PCR duplicates and reads mapping to mitochondrial DNA. Following peak calling, the consensus peak set across all samples was generated. To assess quality metrics we employed functions from the deepTools package^64^ and explored the enrichment of read coverage (plotFingerprint) and visualized gene scores across genome regions (computeMatrix and plotProfile,).

Next, we extracted the peak count matrix and excluded peaks which were present in less than 5 samples. DESeq2 (v1.42.0)^57^ was used to identify differentially accessible peaks. The model was fitted to the count matrix using the DESeqDataSetFromMatrix function. Following statistical tests (Wald test) to compare read counts between the conditions of interest. Peaks with a q-value (Benjamini-Hochberg corrected) below 0.05 were considered differentially accessible (Supplementary Table 4e). Differentially accessible regions were annotated to their genomic regions using ChIPseeker (v1.40.0)^65^ and further analysed using EnrichR^58,59^ and GSEA^60^.

Motif enrichment analysis was conducted on genomic regions that exhibited increased accessibility following IPF or IPF-treatment. The tool findMotifs.pl of the HOMER suite^66^ generated tables listing enriched motifs with statistical significance scores and their associated transcription factors. As a baseline for motif enrichment, the background sequences were randomly selected from the genome. To identify specific instances of the JUNB binding motif, we utilized the tool scanMotifGenomeWide.pl to search for matching positions across the genome.

### Nuclei isolation and scATAC-seq

For scATAC, AOs were singularized. Cells were resuspended in ice-cold buffer (1xPBS + 0.04% BSA) and filtered through a 40µm cell filter (Flowmi® BelArt). Cell concentration, viability, and aggregate were determined with the NucleoCounter NC-3000 (Chemometec) using Via-1 cassette and on NucleoView NC-3000 software (Chemometec, v.2.1). A total of 850 000 cells per sample, with an average cell viability of 92.2 ± 1.7% and cell aggregate <5%, were used for nuclei isolation using 10x Genomics Demonstrated Protocol for Nuclei Isolation for Single Cell Multiome ATAC + GEX Sequencing (CG000365-RevC). Nuclei concentration was determined with the NucleoCounter NC-3000 using A2 slides and a double staining of Acridine Orange and DAPI (Solution 13®, Chemometec). 4 single-nuclei suspensions were employed to generate scATAC libraries which were prepared with the Chromium Next GEM Single Cell Multiome kit ATAC + Gene Expression according to the manufacturer’s instructions (CG000338 Rev E, 10x Genomics). Samples were loaded into a NextGEM Chip J (10x genomics) with a targeted nuclei recovery of 8000 nuclei. Capture and GEM generation was conducted with the Chromium controller followed by reverse transcription, Post-GEM clean-up and a Pre-amplification of 8 cycles. After Pre-Amplification SPRI Cleanup, sample was divided and used as input for scATAC library preparations. For scATAC-seq, 35µL from the purified Pre-Amplification eluate was used for cDNA amplification with 12 cycles and purified with SPRI beads. Qualitative and quantitative measurement of cDNA was conducted with the High Sensitivity NGS Fragment 1-6000bp kit on a 48-channel Fragment Analyzer (Agilent) and 1x dsDNA kit on the Qubit 4 Fluorometer (ThermoFisher), respectively. scATAC-seq libraries were amplified with 12 and 7 PCR cycles and quantified with the High Sensitivity dsDNA Quanti-iT Assay Kit (ThermoFisher) using a Synergy HTX microplate reader (BioTek) and qualitatively assessed with the High Sensitivity NGS Fragment 1-6000bp Kit on a 96-channel Fragment Analyzer (Agilent). Final libraries were normalized, pooled, spiked in with 5% PhiX Control v3 (Illumina) and sequenced on an Illumina Novaseq 6000 at a depth of ∼50,000 reads/cell with dual index, paired end reads (scATAC-seq Read parameters: Rd1: 50bp, Rd2: 8bp, Rd3: 24bp Rd4: 49bp).

### Analysis of scATAC-seq

De-multiplexed scATAC-seq FASTQ files were run in the Cell Ranger ARC pipeline (10X Genomics, v2.0.2) to produce barcoded count matrices of fragment files of ATAC data. GRCh38.86 was used as reference genome for alignment of the reads.

The downstream analysis of the scATAC-seq data was performed with the R package ArchR (v1.0.2)^67^. The basic input for ArchR are Arrow files, a tool-specific data structure that stores data associated with an individual sample (i.e. metadata, accessible fragments, and data matrices). The tab-separated fragment files from the cellranger ARC run were the basis to generate such Arrow files for each of the four input samples. Common quality control metrics such as the number of unique nuclear fragments, the fragment size distribution and transcription starting site enrichment were inspected. Potential doublet cells were filtered out with filterDoublets and the parameter filterRatio set to 1.5.

For dimensionality reduction we used Latent Semantic Indexing (LSI) as suggested in the common workflow due to the high level of sparsity inherent to ATAC-seq data. LSI was calculated via addIterativeLSI, using the “TileMatrix” as input with the following parameters: iterations = 2, resolution = 0.2, sampleCells = 10000, varFeatures = 25000, dimsToUse = 1:30. As we expected sample-specific effects, batch correction was performed with Harmony^68^ based on the LSI space. The 2D embedding was generated on the Harmony-corrected space via addUMAP with the parameters nNeighbors = 40, minDist = 1, metric = “cosine”. We calculated cluster specific markers with getMarkerFeatures and default parameters and assigned cell population identity based on common cell type marker of lung airway epithelial cells (MUC5B, SCGB1A1, KRT5, KRT15). The cell type annotation was used as basis for peak calling with addReproduciblePeakSet which is based on the tool macs2^69^.

We subset to the basal cell population to perform differential accessibility analysis comparing IPF-derived cells to donor cells (getMarkerFeatures with useMatrix = “PeakMatrix”, groupBy = “health_status”, testMethod = “Wilcoxon”, bias = "TSSEnrichment" and "log10(nFrags)") (Supplementary Table 4a).

Differentially accessible regions of basal cells were annotated to their genomic regions and further analysed as described for bulkATAC-seq (Supplementary Table 4b-d). Finally, we aimed to infer which transcription factors may mediate the creation of these accessible chromatin sites. Thus, motif enrichment analysis was performed with peakAnnoEnrichment by providing differentially accessible sites in IPF patients as input (FDR < 0.1 and log2 foldchange >= 0,5).

### Transfections (CRISPR-Cas9/Plasmids)

2D cultured airway epithelial cells or mesenchymal cells were transfected via electroporation with Amaxa Nucleofector 2b device (Lonza) and the respective kit, (Lonza, #VPI-1005 or) at program T-020 according to manufacturer’s instructions to introduce ribonuclear-protein-complex (RNP) complex, plasmids, or control, respectively. The following TrueGuide Synthetic gRNAs (Thermo Fisher, #A35533) were used: OGT (CRISPR887744_SGM); JUNB (CRISPR924803_SGM, CRISPR924804_SGM ratio 1:1 mixed); TrueGuide sgRNA Negative Control (#A35526).

### AAV6.2 production

Self-complementary AAV6.2 vectors were produced and purified as previously described^70^. Briefly, high-density HEK-293h cells (AcCELLerate) were seeded into 8-layer CELLdiscs and triple-transfected with pHelper, pRep2/Cap6.2 and JUNB-encoding plasmids (Supplementary Table 12) using the Calcium phosphate procedure. Following culture for 72 h, cells were lysed by three freeze/thaw cycles under high salt conditions. Following salt-active nuclease treatment and PEG precipitation, AAVs were purified by iodixanol step gradient ultracentrifugation, followed by Amicon-15-based buffer exchange/ultrafiltration and sterile filtration. Genomic titers were determined by dPCR (Qiagen), using ITR-specific primers.

AOs were infected after seeding with AAV constructs for 24 h at a MOI of 10^5^.

### Rat precision cut lung slices (rPCLS)

Male rats (Wistar Han) were purchased from (Charles River). Rats were housed in individually ventilated cages at 22–25°C, a humidity of 46–65%, and 12-h day/night cycle. Animals received water and food ad libitum. Ethical approval for this study was obtained from the regional govern-mental animal care and use office (Regierungspräsidium Tübingen, Germany, TVV 18-030-O).

Healthy rats were euthanized using 60 mg/kg Pentobarbital-sodium (Narcoren). Diaphragm was punctured and lungs were infiltrated with around 20 ml of LM agarose using a cannula with syringe. Immediately, agarose-filled lungs were cooled with ice for 15 min. Afterwards, ice was removed, the lobes were separated and cut with a microtome (Krumdieck) to a thickness of 200-350 µm on medium blade and movement speed. Once slicing is completed, slices were transferred to 6-well plate inserts with 10 ml medium (1x DMEM/F12 + L-glutamine + 15 mM HEPES + 1x Antibiotica-Antimycotica + 0.1 % fetal bovine serum) and cultivated at 37 °C, 5 % CO2. The next day, slices were distributed in a 48 well plate with 300 µl medium including respective treatment. After 48 h, medium was exchanged and slices cultivated another 72 h for a total cultivation time of 144 h.

### RNA and protein extraction of rPCLS

Slices were collected in Precellys Lysis Matrix D tubes and snap-frozen in liquid nitrogen. Afterwards, protein was lysed in T-Per (+phosphatase/protease inhibitor) buffer, RNA was lysed using RLT Plus buffer and samples were homogenized for 25 s at 6500 rpm on a Precellys device (Bertin Technologies).

For protein, samples were incubated on ice for 15 min and again homogenized with the previous settings. Next, samples were cleared at 18,000 g for 15 min, followed by simple western analysis.

For RNA, homogenates were incubated for 5 min at RT and transferred to PhaseLock tubes with phenol/chloroform/isoamyl alcohol (25:24:1). After shaking, samples were centrifuged at 16,000g for 5 min and chloroform/isoamyl alcohol (24:1) was added followed again by shaking. Next, samples were incubated for 3 min at RT and centrifuged again for 10 min. The extracted RNA was then purified and processed for RT-PCR as previously described.

### Statistical analysis and reproducibility

GraphPad prism (version 10.1.2) was used to perform statistical analysis. The following tests were used to determine significance: Unpaired t-test, paired t-test with Wilcoxon test, one-way ANOVA with Holm-Šídák’s, Šídák’s, Tukey’s or Dunn’s test, non-linear/linear regression. Gaussian distribution was determined for all tests, which required normal distribution, with the Kolmogorov-Smirnov test (α = 0.05). At least three biological replicates were performed for all experiments.

## Acknowledgements

We thank A. Arduini for critical reading of the manuscript, D. Jonigk and the German Centre for Lung Research (DZL/BREATH) Hannover for providing us with patient-derived lung tissue, I. Kollak, E. Peter, J. Hill, D. Schäfer, S. Klee, L. Di Niro, R. Ammann, B. Lämmle and the Drug Discovery Sciences department for technical assistance and support. This work was funded by Boehringer Ingelheim Pharma GmbH und Co. KG. All drawings were created using Biorender.com.

## Author contributions

MB designed, performed, and supervised most of the experiments and analysed data. DS, MB, WR, AD performed sequencing experiments. MA, FR analysed sequencing data. ZA, LH, VS performed experiments. MB and JB performed *ex vivo* experiments. MJT, ML, HS, FG, BS, CV provided conceptual advice. HQL conceived, conceptualized, supervised the study, and designed experiments. MB and HQL wrote the paper. All authors commented on and edited the manuscript.

## Data availability

ATAC- and RNA-seq data generated within this study was deposited in the Gene Expression Omnibus and will be made accessible upon acceptance of the manuscript. All other data is available on request.

**Extended Figure 1: Patient-derived AOs recapitulate important fibrotic characteristics.**

**a**, RT-PCR analysis of airway-specific markers *KRT5*, *SCGB1A1*, *MUC5B*, and *FOXJ1* were obtained from n = 5 control AOs (mean + s.e.m) at 2 and 3 weeks of cultivation. Air-liquid interface (ALI) differentiated airway cells serve as positive, while submerged-cultivated airway cells as negative control.

**b**, Representative H&E staining of IPF and control AOs after 21 days showing lumen and cilia. **c**, Representative PAS staining of control AOs after 21 days, confirming glycolytic mucus presence within organoids.

**d**, Bright-field images of AOs on day 0 and 15 post-seeding, revealing increased size in IPF AOs.

**e**, Growth rate constant (k) determination from n = 4 IPF and n = 7 control AO donors over 15 days, indicating significantly increased proliferation in IPF AO (mean + s.d, t-test, **p < 0.01). **f**, Nanoindentation measurement shows increased viscoelasticity (Young modulus) in IPF AOs from n = 5 IPF and n = 5 control donors (boxplot, ****p < 0.0001).

**g**, Increased glucose capacity in IPF AOs (n = 5, mean + s.e.m., \**p* < 0.05, ANOVA/ Tukey’s).

**Extended Figure 2: Aberrant OGlcNAcylation is involved in IPF.**

**a**, Schematic representation of the experimental procedure. Singularised AOs were seeded and cultivated for 4 days to induce organoid formation and grown for another 7 days. Additionally, they could be exposed to IPF stimuli for 7 days to increase the fibrotic phenotype. After a total of 11 days, AOs were collected and processed for multiome bulk RNA-/ATAC-seq or scATAC-seq.

**b**, PCA of RNA-seq data with the top 1000 variable genes (*q-*value < 0.05) according to coefficient of variation shows distinct clustering of IPF and control AOs.

**c**, Hierarchical clustering of top 50 DEGs from RNA-seq comparing control and IPF AOs including treatment with IPF-stimuli (*q*-value < 0.05).

**d**, Reactome analysis of IPF-stimulated control AOs.

**e**, Reactome analysis of IPF-stimulated IPF AOs.

**Extended Figure 3: O-GlcNAcylation plays an important regulatory role in IPF.**

**a**, **b**, RT-PCR shows that OGT deletion in airway epithelial cells (**a**) and lung fibroblasts (**b**) leads to decrease in *OGT* expression after IPF stimulation (n = 6, mean + s.e.m, \*\*\*\**p* < 0.0001, ANOVA/ Tukey’s).

**c**, **d**, RT-PCR reveals attenuated expression of profibrotic genes *COL1A1* (**c**) and *FN1* (**d**) upon OGT deletion in fibroblasts after fibrotic induction with IPF-stimuli (n = 6, mean + s.e.m, \*\*\**p* < 0.001, \*\*\*\**p* < 0.0001, ANOVA/ Tukey’s).

**e**, **f**, Determination of EC_50_ values for OSMI-4 in AOs based on relative O-GlcNAc expression from western blot analysis (**e**) and quantification (**f**) of AO protein lysates treated within a range of 1.25-20 µM OSMI-4 for 7 days. DMSO was used as negative control.

**g**, **h**, Determination of EC_50_ values for OSMI-4 in airway epithelial cells based on relative O-GlcNAc expression from western blot analysis (**g**) and quantification (**h**) of AO protein lysates treated within a range of 1.0-40 µM OSMI-4 for 72 h. DMSO was used as negative control.

**i**, Schematic overview of epithelial-mesenchymal-coculture set up and treatment strategy. Airway epithelial cells were grown for 10 days in air-liquid-interface, before being exposed to IPF-stimuli for another 14 days to induce chronic injury. Afterwards, injured airway epithelial cells (tSAECs) and lung fibroblasts were cocultured for another 72 h.

**Extended Figure 4: JUNB TF motif enrichment in IPF basal cells promotes fibrotic cell fate.**

**a**, **b**, UMAP representation of single-cell chromatin accessibility (scATAC-seq) clustering in IPF and control AOs (**a**) and IPF cell types (**b**) (basal, proliferating basal, secretory and aberrant basaloid cells).

**c**, PCA of top 5000 variable regions (bulk ATAC-seq) (*q*-value < 0.05) according to coefficient of variation shows distinct clustering of IPF and control-derived AOs with and without IPF-RC treatment.

**d**, **e**, Genomic annotations of IPF AOs (bulk ATAC-seq) open (**d**) and closed (**e**) differential accessible regions (DARs, *q*-value < 0.05) reveals enriched open chromatin in IPF promotor regions close to the transcription start site (TSS).

**f**, GO enrichment analysis of protein coding, open DARs (*q*-value < 0.05) in IPF AOs (bulk ATAC-seq) shows involvement of developmental and proliferative pathways.

**g**, Bioplanet analysis of closed DARs (*q*-value < 0.05) in IPF AOs (bulk ATAC-seq) shows enrichment of metabolic pathways.

**h**, Transcription factor (TF) motif enrichment analysis shows upregulation of different activator protein 1 (AP-1) motifs in IPF basal cells (scATAC-seq).

**i - k**, TF motif enrichment analysis (bulk ATAC-seq) in IPF vs. Control AOs (**i**), Control AOs treated with IPF-stimuli vs. CTRL (**j**), and IPF AOs treated with IPF-stimuli vs. CTRL (**k**) reveals increased motif enrichment for AP-1 family members in IPF conditions.

**l**, RT-PCR shows decreased *JUNB* expression upon JUNB deletion in airway epithelial cells after fibrotic induction with IPF-stimuli (n = 5, mean + s.e.m, ANOVA/Tukey’s).

**m,** RT-PCR reveals attenuated expression of profibrotic genes (i.e. *COL5A1*, *FN1*) upon JUNB deletion in lung fibroblasts after IPF-stimuli treatment (n = 6, mean + s.e.m, p* < 0.05, p*** < 0.001, ANOVA/Tukey’s).

**Extended Figure 5: O-GlcNAcylation of JUNB is involved in the pro-fibrotic response.**

**a,** Representative simple western analysis of O-GlcNAc immunoprecipitated fraction showing increase of O-GlcNAc mark on JUNB in injured n = 3 lung fibroblast donors. IgG antibody was used as negative control.

**b,** RT-PCR reveals increased expression levels of JUNB after JUNB-LOF induction deletion (n = 6, mean + s.e.m, *p < 0.05). Deletion of JUNB was employed as controls.

**c, d,** RT-PCR showcases diminished expression of *COL1A1* (**c**) and *MMP10* (**d**) after LOF-induction and injury with IPF-stimuli in airway epithelial cells (n = 6, mean + s.e.m, *p < 0.05, **p < 0.01, ***p < 0.001, ****p < 0.0001, ANOVA/ Šídák’s) comparable to the effect of JUNB-KD (n = 2 mean + s.e.m). Plasmid was generated by site-specific mutations in the O-GlcNAc sites of JUNB. Expression levels of *JUNB* (**b**) increased upon induction of the JUNB-LOF plasmid and decreased in the KD.

**e**, **f**, RT-PCR shows reduced fibrotic gene expression upon JUNB-LOF transfection and fibrotic treatment with IPF-stimuli representatively shown for *COL1A1* (**e**) and *FN1* (**f**) in lung fibroblasts (n = 7, mean + s.e.m, **p < 0.01, ***p < 0.001, ****p < 0.0001, ANOVA/ Tukey’s). **g**, RT-PCR reveals increased *JUNB* levels upon induction of JUNB-LOF in lung fibroblasts (n = 7, mean + s.e.m,, ****p < 0.0001, ANOVA/ Tukey’s).

**Extended Figure 6: OSMI-4 inhibits O-GlcNAc via blocking OGT in rPCLS.**

**a**, Representative simple western analysis of O-GlcNAc fraction in rPCLS treated with +/- r-IPF-RC and +/- OSMI-4 showed increase of O-GlcNAc mark after stimulation with r-IPF-RC for 120 h and decreased O-GlcNAc after addition of OSMI-4 for 120 h or 72 h.

